# Alignment-free sequence composition based features can distinguish paralogous proteins

**DOI:** 10.1101/2024.04.26.591291

**Authors:** Pranav Machingal, Rakesh Busi, Nandyala Hemachandra, Petety V Balaji

## Abstract

Paralogous proteins have a common ancestor but have diverged in functionality. Homology- and similarity-based methods often fail to distinguish paralogous proteins, due to a nuanced sequence-function relationship that remains largely intractable. Going beyond these methods, we extract various sequence composition based feature representations from proteins and use the same to train classifiers for distinguishing paralog pairs. We consider 20 paralog pair datasets and train a linear support vector machine (SVM) classifier for each dataset to distinguish among them. Amino acid composition (AAC) and 2- ordered amino acid composition (2-OAAC) features achieve high predictive performance across these datasets, despite the low sample size and potential label noise. Furthermore, we utilise the marginal contribution feature importance (MCI) to identify specific amino acids driving classification. We show that retaining only the top-*k* MCI-ranked amino acids preserves nearly all predictive power while acting as a regularizer. Structural mapping of these top-ranked features in families such as globins, tubulins, and histones reveals that they frequently localise to critical functional and oligomerisation interfaces. Ultimately, our interpretable machine learning framework successfully bridges sequence composition and protein activity, providing a scalable computational method for identifying potential functional determinants that can guide future wet-lab site-directed mutagenesis experiments to study protein function. The code and data are available at: https://github.com/Pranav-Machingal/paralog-classification

## 1. Introduction

Proteins form the fundamental machinery in living systems, having several vital functions such as DNA replication, catalysis, transport, environmental interaction, etc. Advancements in sequencing technologies have resulted in exponential growth of protein sequence databases.^1^ However, the number of experimentally verified annotations constitutes a tiny fraction: only 0.57% of 250 million sequences in UniProtKB^1^ have manually reviewed annotations. Experimental methods for determining biological process level functions (transcription, DNA repair, etc.) are high-throughput, whereas methods for molecular function (catalysis, ligand specificity, etc.) are low-throughput and hence are not scalable. The relationship between sequence and function is subtle and has not yet been fully decoded.

Paralogs are proteins that have a common ancestor but have diverged functionally. The functional difference in two paralogous proteins is considered to arise due to evolutionary changes in the sequences.^2^ As a result, paralogous proteins often share striking similarities in their primary sequences and tertiary structures, yet exhibit critical differences in binding specificity, enzymatic activity, cellular localisation, or regulatory roles. Traditional homology-based sequence alignment and global similarity metrics frequently struggle to differentiate between many paralog pairs.^3^ As illustrated in Fig. 1, for two protein families that are paralogous, the within-class and inter-class sequence similarities are, in many cases, indistinguishable (e.g., Trypsin vs Chymotrypsin, LDHs: A vs B, GPCRs: Lipid vs Peptide). A simple sequence similarity cutoff cannot distinguish the paralog pairs in these cases.

**Fig. 1.**
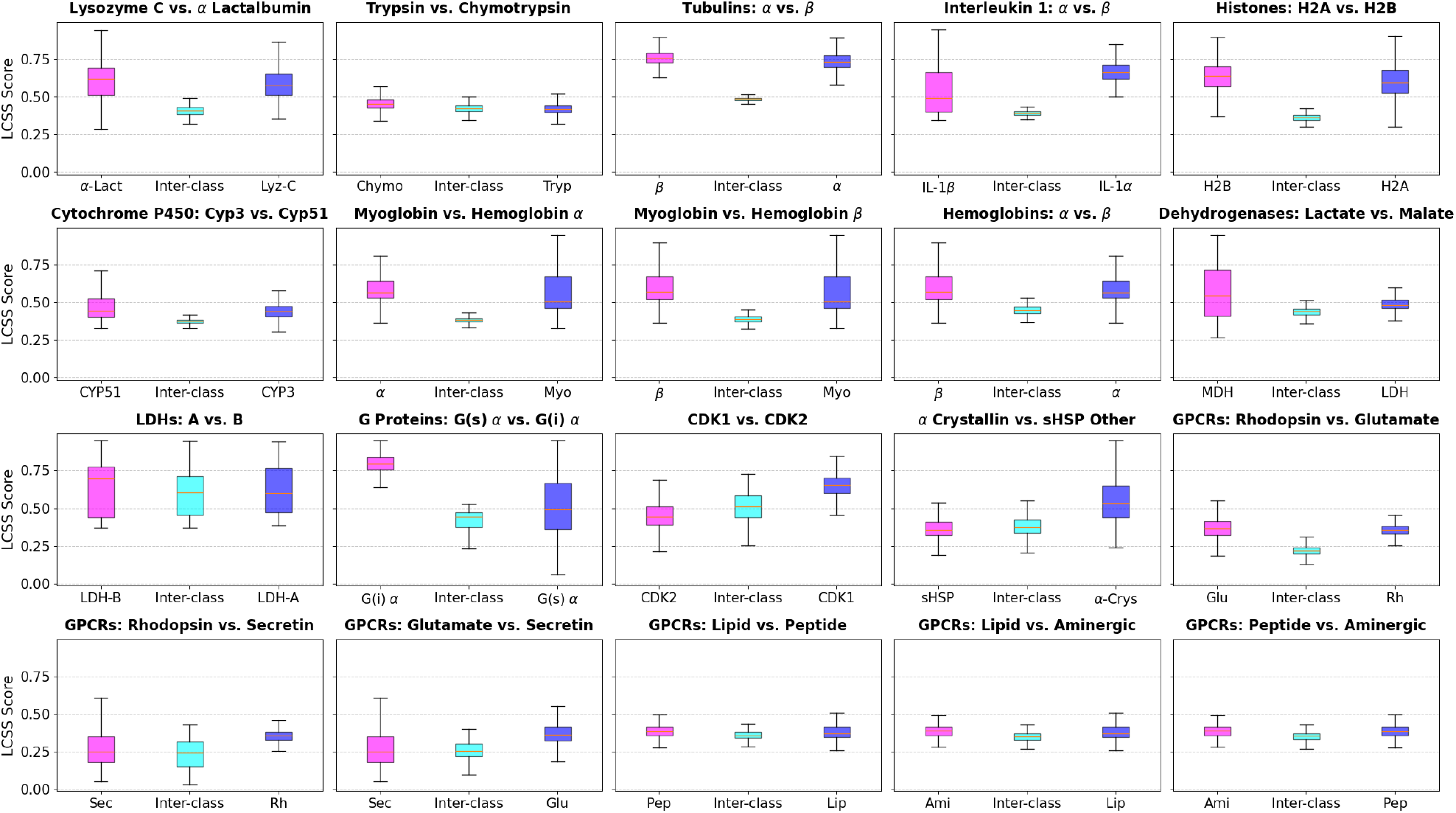
Within-class and inter-class longest common subsequence score (LCSS) for paralog pairs. LCSS between two sequences is defined as the length of their longest common (non-contiguous) subsequence divided by the length of the longer of the two sequences.

In this work, we cast the problem of differentiating highly homologous protein families as a classification task across diverse paralog pairs. Rather than relying on standard sequence alignment, we transform the primary sequences into comprehensive feature spaces based on sequence composition. We utilise both fundamental amino acid composition (AAC) and higher-order representations such as 2-ordered amino acid composition (2-OAAC), alongside physicochemical properties-based groupings (e.g., Dayhoff, Taylor, and Conjoint Triad classifications). By training linear support vector machine (SVM) classifiers on these representations, we demonstrate that compositional features are often sufficient to achieve high predictive accuracy. We consider a diverse dataset of 20 paralog pairs and train linear SVMs to distinguish each pair.

Applying data-driven machine learning (ML) models to differentiate these paralogs introduces its own set of data-centric challenges. Primarily, there is a stark trade-off between sequence quality and volume: relying exclusively on manually reviewed sequences (from SwissProt^1^) typically yields extremely small sample sizes for many specific paralog families, which severely limits the generalisation of ML models, especially when using high-dimensional informative features. Conversely, expanding the dataset to include unreviewed, computationally annotated entries (from TrEMBL^1^) introduces significant noise, such as misannotations (label noise). Consequently, any successful ML model must be highly resilient to noisy, sparse data while still extracting meaningful biological signals. Details of our data collection and pre-processing are provided in Section 2.1.

A typical experiment to investigate the role of an (or a group of) amino acid(s) in the function of a protein is to perform a site-directed mutagenesis experiment: replace one or more amino acids and test the effect of the sequence change.^4^ While wet-lab experimental mutagenesis is the gold standard for identifying functional determinants, it is inherently low-throughput and challenging to scale. Crucially, our objective extends beyond mere classification; we aim to rigorously interpret the biological determinants driving these predictions to generate testable hypotheses. We utilise marginal contribution feature importance (MCI)^5^ to evaluate the characteristic contribution of the different amino acid types in each of the paralog pair classification tasks. This allows us to identify a subset of amino acids that potentially contribute to the functional divergence of the paralog pairs, providing a hypothesis for site-directed mutagenesis experiments to study protein function.

We discuss our methodology and computational results in detail in the next sections.

## 2. Datasets, feature engineering, classifier design and feature importance

We discuss the following aspects of our methodology in this section: datasets, feature engineering, classifier design and feature importance

### 2.1. Paralog pair datasets

We consider a diverse dataset of 20 paralog pairs, comprising 31 protein families (and subfamilies). These are listed in Table 1. The paralog pairs were selected based on the availability of sufficient sequences to train and evaluate the paralog-pair classifiers. All the sequences were collected from UniProtKB^1^ (both reviewed and unreviewed), and the specific queries used for collecting the sequences are available in Supplementary File 1. The datasets show a range of sequence and functional diversity. We use the longest common subsequence score (LCSS) metric to quantify sequence diversity, see Fig. 1. The dataset contains paralog pairs with inter-class LCSS ranging from very low (e.g., GPCRs) to very high (e.g., LDHs). Functional diversity, as discerned from biology literature, also shows a wide range, from subtle functional differences (e.g., trypsin/chymotrypsin) to drastic ones (e.g., lysozyme c/*α*-lactalbumin).

**Table 1.** Summary of the protein families datasets used in this study. The train/test splits are 80:20.

| Family | Train | Test | Organisms |
| --- | --- | --- | --- |
| **Lysozyme-like** |  |  |  |
| Lysozyme C | 54 | 14 | 51 |
| α-Lactalbumin | 67 | 17 | 74 |
| **Trypsin-like protease** |  |  |  |
| Trypsin | 1683 | 421 | 388 |
| Chymotrypsin | 180 | 46 | 80 |
| **Tubulin** |  |  |  |
| Tubulin $\alpha$ | 64 | 16 | 60 |
| Tubulin $\beta$ | 118 | 30 | 125 |
| **Interleukin-1 (IL-1)** |  |  |  |
| IL-1 $\alpha$ | 43 | 11 | 52 |
| IL-1 $\beta$ | 79 | 20 | 92 |
| **Histone** |  |  |  |
| H2A | 230 | 58 | 175 |
| H2B | 87 | 22 | 72 |
| **Cytochrome P450** |  |  |  |
| Cyp3 | 298 | 75 | 102 |
| Cyp51 | 194 | 49 | 202 |
| **Globins** |  |  |  |
| Myoglobin | 205 | 52 | 215 |
| Hemoglobin-α | 267 | 67 | 223 |
| Hemoglobin-β | 224 | 56 | 205 |
| **GPCRs** |  |  |  |
| Rhodopsin-like | 7808 | 1952 | 646 |
| ↳ Lipid | 298 | 75 | 164 |
| Peptide | 1986 | 497 | 395 |
| Aminergic | 2782 | 696 | 439 |
| Glutamate-like | 360 | 91 | 186 |
| Secretin-like | 2946 | 737 | 429 |
| **Dehydrogenases** |  |  |  |
| Lactate | 2594 | 649 | 2813 |
| ↳ LDH-A (muscle-type) | 52 | 13 | 59 |
| LDH-B (heart-type) | 57 | 15 | 65 |
| Malate | 950 | 238 | 1076 |
| **G proteins** |  |  |  |
| G(s) $\alpha$ | 591 | 148 | 421 |
| G(i) $\alpha$ | 45 | 12 | 52 |
| **Cyclin-dependent kinases (CDKs)** |  |  |  |
| CDK1 | 551 | 138 | 645 |
| CDK2 | 452 | 114 | 217 |
| **Small heat shock proteins (sHSPs)** |  |  |  |
| α-Crystallin | 73 | 19 | 76 |
| sHSP Other | 108 | 28 | 66 |

A sequence redundancy filter with a threshold of 0.95 is used to remove highly similar sequences from the dataset. The sequences collected from each family are randomly split 80:20 into training and test sets, which are then used to train and evaluate the paralog pair classifiers.

### 2.2. Sequence composition based feature engineering

To distinguish between the highly homologous paralog pairs without relying on full sequence alignment, we map the raw protein sequences into various lower-dimensional feature spaces based on their composition. We consider 7 purely composition-based features that use no sequence order information. And one feature that uses composition with partial sequence order information. These are discussed below

- Composition only features:
  a. **(AAC):** A 20-dimensional vector containing the normalised occurrence frequencies of the standard 20 amino acids.
  b. **4-group composition:** A 4-dimensional vector based on side-chain properties: nonpolar (A, V, L, I, P, F, W, M), polar (G, S, T, C, Y, N, Q), positively charged (K, R, H), and negatively charged (D, E).
  c. **6-group chemical composition:** A 6-dimensional representation classifying amino acids by their structural functional groups: aliphatic (G, A, V, L, I, P), aromatic (F, Y, W), sulfur-containing (C, M), hydroxylated (S, T, Y), acidic & amides (D, E, N, Q), and basic (K, R, H).
  d. **Dayhoff’s composition:** A 6-dimensional feature vector based on Dayhoff’s physical-chemical groups:^6^ sulfhydryl (C), small (S, T, P, A, G), acidic & amide (N, D, E, Q), basic (H, R, K), aliphatic (M, I, L, V), and aromatic (F, Y, W).
  e. **Conjoint triad:** A 7-dimensional vector grouping amino acids by their dipole scales and side-chain volumes:^7^ group 1 (A, G, V), group 2 (I, L, F, P), group 3 (Y, M, T, S), group 4 (H, N, Q, W), group 5 (R, K), group 6 (D, E), and group 7 (C).
  f. **Taylor’s composition:** A 9-dimensional vector based on Taylor’s overlapping Venn diagram classification:^8^ hydrophobic (V, I, L, F, Y, W, M, A, C, T, H, K), polar (S, T, Y, N, D, Q, E, K, R, H, W), small (P, A, G, C, S, V, N, D), tiny (A, G, S), aromatic (F, Y, W, H), aliphatic (I, L, V), charged (D, E, K, R, H), positive (K, R, H), and negative (D, E).
  g. **Multilabel property encoding:** A 4-dimensional feature vector based on interaction capabilities of amino acids: hydrogen bond donor (S, T, Y, N, Q, W, K, R, H), hydrogen bond acceptor (D, E, N, Q, S, T, Y, H), *π*-Stacking capable (F, Y, W, H), and disulfide bond capable (C).

The features other than AAC discussed above can be seen as low-dimensional (4-9 dimensions) linear projections of the AAC features. These features are based on various ways of grouping amino acids by their physicochemical properties.

- Feature using partial sequence order information:
  a. 2**-ordered amino acid composition (**2**-OAAC):** The 2-OAAC feature tracks the normalised occurrences of ordered 2-tuples of amino acids appearing as subsequences (not necessarily contiguous) in the protein sequence. This results in a 400-dimensional (20^2^) feature vector. A length *L* sequence will have 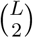 possible subsequences of size 2. For a protein sequence **p** = (*p*_1_, *p*_2_, …, *p*_*L*_) of length *L* with *p*_*j*_ ∈ *A* being one of the standard 20 amino acids, the 2OAAC feature **x**^2*OAAC*^ ∈ ℝ^400^ for **p** is computed as follows,

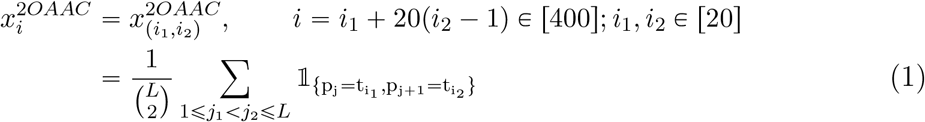

With 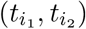 ∈ *A* ×*A* being an order 2-tuple of amino acid types. 2-OAAC can be generalised to *k*-OAAC by considering ordered *k*-tuples, and thereby encoding higher orders of sequence order information for higher *k*. However, this comes at the cost of an exponentially large number of features (20^*k*^). For *k* =1, it is the same as AAC.

### 2.3. Linear SVM Classifiers

Using each of the features discussed in Section 2.2, we trained linear support vector machine (SVM) classifiers to distinguish between the two families from each paralog pair in our dataset. We use the `LinearSVC` implementation from the `scikit-learn` library. To address potential class imbalances, the models were optimised using a class-balanced hinge loss with a maximum of 50,000 iterations. The class-balanced hinge loss has been shown to be statistically consistent with the AM score, the arithmetic mean of class-wise recall.^9^ The AM score, also known as balanced accuracy,^10^ is a popular metric for binary classification with imbalanced datasets.

To select the optimal regularisation hyperparameter *C*, we performed a two-stage coarse-to-fine grid search using 10-fold cross-validation on the training data, optimising for the AM score. The coarse search scanned a logarithmic grid ranging from 10^−3^ to 10^3^. A subsequent fine search was then conducted across 11 logarithmically spaced points centred around the best coarse value.

We report the AM scores on the test data for all the trained classifiers in Section 3.1.

### 2.4. Marginal contribution feature importance (MCI) scores

Marginal Contribution feature Importance (MCI) is an axiomatic metric for evaluating feature importance.^5^ For a feature set *T*, the MCI score of a feature *i* ∈ *T* is defined as its maximum marginal contribution to any subset *S* ⊆ *T\*{*i*}:

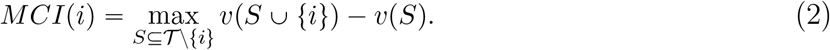

Here, the value function *v*(*S*) = 1 − *tr_er*(*S*) measures the linear separability of the classes using only the features in *S*.^11^ We compute the class-balanced hinge loss *tr_er*(*S*) via the linear SVM objective:

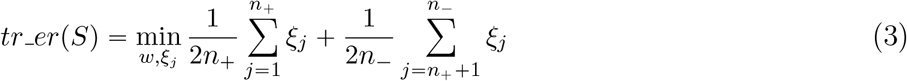

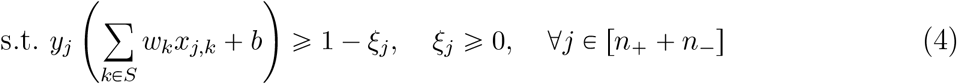

where *n*_*+*_ and *n*_−_are the number of positive and negative samples, and {(*x*_*j*_, *y*_*j*_)} is the training data. The loss *tr_er*(*S*) ranges from 0 (perfect linear separability with *S*) to 1 (for *S* = Ø).

Exact MCI computation requires evaluating 2^|*T* |^ subsets, so we use a linear-time Monte Carlo sampling approximation.^12^ However, the variance of this approximation increases significantly with |*T* |. For high-dimensional features like 2-OAAC (|*T* | = 400), computing feature-wise MCI yields unstable rankings. Instead, we devise an amino acid-wise MCI scoring strategy, which is discussed next.

#### Amino acid-wise MCI for 2-OAAC features

We can view the 400-dimensional 2-OAAC features as a 20 × 20 matrix corresponding to ordered pairs of the 20 standard amino acids, denoted by *A*. We define *g*p*S*q as the set of all 2-OAAC features formed exclusively by amino acids in a subset *S* ⊆ *A*. The amino acid-wise MCI for amino acid *i* ∈ *A* is then:

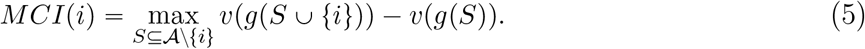

This approach evaluates the importance of the 20 underlying amino acids rather than the 400 individual 2-OAAC features, providing much more stable estimates.

We select feature subsets with high MCI scores and train and evaluate the paralog pair classification task using only these subsets. We do this for AAC and 2-OAAC features. The test scores using different subset sizes are reported in Section 3.1. For the AAC feature we also highlight the role of amino acids with high MCI scores in distinguishing paralogs, in Section 3.2.

## 3. Classification results and MCI analysis

### 3.1. Classification results

The training, validation and test AM scores for 20 paralog pair classifications, using the 8 types of sequence composition based feature engineerings discussed in Section 2.2, are shown in

Fig. 2. Overall, linear SVMs achieve remarkably high test AM scores across most paralog pairs. Also, for most paralog pairs and feature engineering, the tight clustering of training, validation, and test scores indicates that the models generalise well and are robust to overfitting. While simpler features like AAC perform well on many pairs, higher-order representations like 2-OAAC consistently provide better separation for challenging pairs, such as LDHs A vs B (improving from 0.57 to 0.86) and Trypsin vs Chymotrypsin (improving from 0.79 to 0.90). The other composition-only features based on amino acid groupings have either lower or comparable classification performances to AAC/2OAAC across the paralog pairs (for LDHs: A vs B, group composition features worked better than AAC). AAC has test AM scores ⩾ 0.82 for 18 of 20 paralog pairs, ⩾ 0.9 for 13 pairs and ⩾ 0.95 for 11 pairs. While 2-OAAC has test AM scores ⩾ 0.86 for all 20 paralog pairs, ⩾ 0.9 for 19 pairs (0.86 for LDHs: A vs B) and ⩾ 0.95 for 16 pairs.

**Fig. 2.**
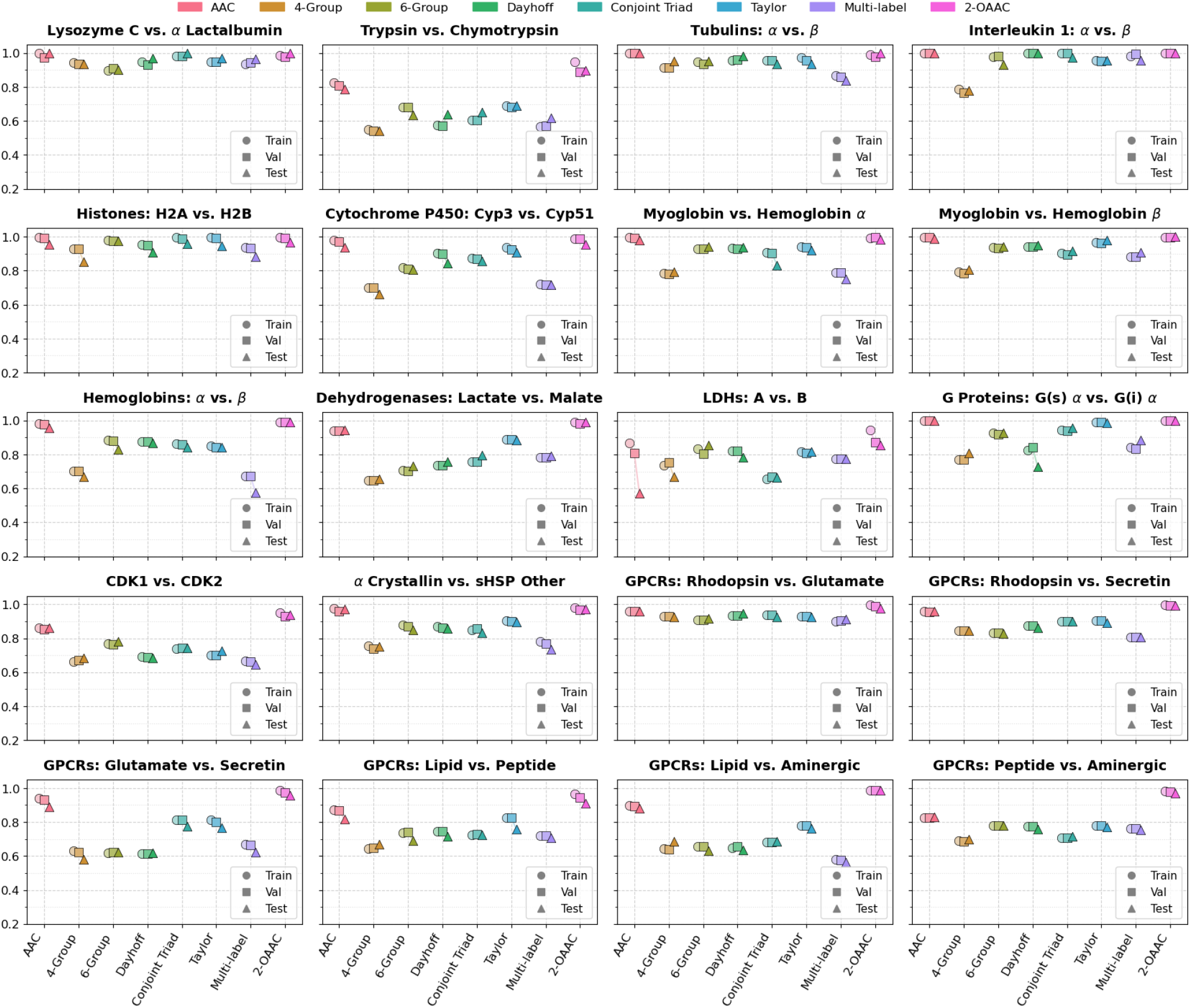
Classification performances across paralog pairs for different sequence composition based feature engineerings. The y-axis shows the AM scores (balanced accuracy).

To test if the information regarding only a subset of amino acids could still distinguish paralog pairs, we re-evaluated classification performance using only the top-*k* amino acids ranked by their MCI scores for AAC and 2-OAAC (Table 2). Remarkably, restricting the feature space to just the top 10 or 7 amino acids preserves nearly all predictive power compared to using the full feature sets. In certain cases, such as the AAC features for LDHs A vs B, the top-*k* subset notably outperforms the full feature set (Test AM increasing from 0.57 to 0.74 for *k* = 10). This suggests that discarding uninformative amino acids acts as an effective regularizer, mitigating noise and reaffirms that the MCI algorithm successfully isolates specific residues potentially driving paralog divergence.

**Table 2.** Classification performance across paralog pairs using full versus MCI-selected features. The AM scores (balanced accuracy) on the test datasets are reported in the table. For 2-OAAC, feature selection is based on the amino acid-wise MCI (Section 2.4), meaning a top-*k* selection yields *k*^2^ features from the 2-OAAC feature vectors. For each feature type and paralog pair, the highest and second-highest scores are highlighted in bold and underlined, respectively.

| Paralog Pair | AAC |  |  |  | 2-OAAC |  |  |  |
| --- | --- | --- | --- | --- | --- | --- | --- | --- |
| | Full | $k = 10$ | $k = 7$ | $k = 5$ | Full | $k = 10$ | $k = 7$ | $k = 5$ |
| Lysozyme C vs α-Lactalbumin | **1.00** | <u>0.97</u> | **1.00** | <u>0.97</u> | **1.00** | <u>0.97</u> | <u>0.97</u> | <u>0.97</u> |
| Trypsin vs Chymotrypsin | **0.79** | <u>0.78</u> | 0.75 | 0.75 | **0.90** | <u>0.84</u> | 0.78 | 0.75 |
| Tubulins: $\alpha$ vs $\beta$ | **1.00** | **1.00** | **1.00** | **1.00** | **1.00** | **1.00** | **1.00** | **1.00** |
| Interleukin-1: $\alpha$ vs $\beta$ | **1.00** | **1.00** | **1.00** | **1.00** | **1.00** | **1.00** | **1.00** | <u>0.98</u> |
| Histones: H2A vs H2B | <u>0.95</u> | **0.97** | **0.97** | **0.97** | <u>0.97</u> | 0.95 | **1.00** | 0.96 |
| Cytochrome P450: Cyp3 vs Cyp51 | 0.94 | <u>0.96</u> | <u>0.96</u> | **0.98** | 0.96 | **0.98** | <u>0.97</u> | 0.96 |
| Myoglobin vs Hemoglobin $\alpha$ | <u>0.98</u> | **0.99** | 0.97 | 0.94 | **0.98** | **0.98** | **0.98** | <u>0.97</u> |
| Myoglobin vs Hemoglobin $\beta$ | **0.99** | 0.97 | <u>0.98</u> | 0.96 | **1.00** | <u>0.98</u> | <u>0.98</u> | <u>0.98</u> |
| Hemoglobins: $\alpha$ vs $\beta$ | **0.96** | <u>0.95</u> | **0.96** | <u>0.95</u> | <u>0.99</u> | **1.00** | <u>0.99</u> | 0.98 |
| Dehydrogenases: Lactate vs Malate | **0.94** | <u>0.93</u> | 0.92 | 0.92 | **0.99** | <u>0.96</u> | 0.93 | 0.91 |
| LDHs: A vs B | 0.57 | **0.74** | <u>0.71</u> | 0.67 | **0.86** | 0.71 | <u>0.72</u> | <u>0.72</u> |
| G Proteins: G(s) $\alpha$ vs G(i) $\alpha$ | **1.00** | <u>0.99</u> | **1.00** | <u>0.99</u> | **1.00** | **1.00** | **1.00** | <u>0.99</u> |
| CDK1 vs CDK2 | <u>0.86</u> | **0.87** | 0.85 | <u>0.86</u> | **0.94** | <u>0.89</u> | 0.85 | 0.81 |
| α-Crystallin vs sHSP Other | **0.97** | **0.97** | <u>0.91</u> | 0.87 | **0.97** | <u>0.95</u> | 0.89 | 0.94 |
| GPCRs: Rhodopsin vs Glutamate | **0.96** | <u>0.95</u> | 0.94 | 0.93 | **0.98** | 0.96 | <u>0.97</u> | 0.92 |
| GPCRs: Rhodopsin vs Secretin | **0.96** | <u>0.93</u> | 0.90 | 0.89 | **0.99** | <u>0.98</u> | 0.97 | 0.92 |
| GPCRs: Glutamate vs Secretin | **0.89** | **0.89** | **0.89** | <u>0.78</u> | **0.96** | 0.91 | <u>0.92</u> | 0.88 |
| GPCRs: Lipid vs Peptide | **0.82** | **0.82** | <u>0.81</u> | 0.75 | **0.91** | <u>0.88</u> | 0.86 | 0.81 |
| GPCRs: Lipid vs Aminergic | **0.88** | 0.85 | <u>0.87</u> | 0.86 | **0.99** | <u>0.98</u> | 0.97 | 0.97 |
| GPCRs: Peptide vs Aminergic | **0.83** | 0.77 | <u>0.78</u> | 0.75 | **0.97** | <u>0.92</u> | 0.90 | 0.86 |

We also see that paralog pairs having the highest MCI score for AAC features < 0.4 (See Fig. 3, trypsin/chymotrypsin, LDH-A/LDH-B, CDK1/CDK2 and some of the GPCR pairs), have < 0.9 test AM scores with full AAC features. However, a significant jump in the test AM scores is observed for these pairs with 2-OAAC features.

**Fig. 3.**
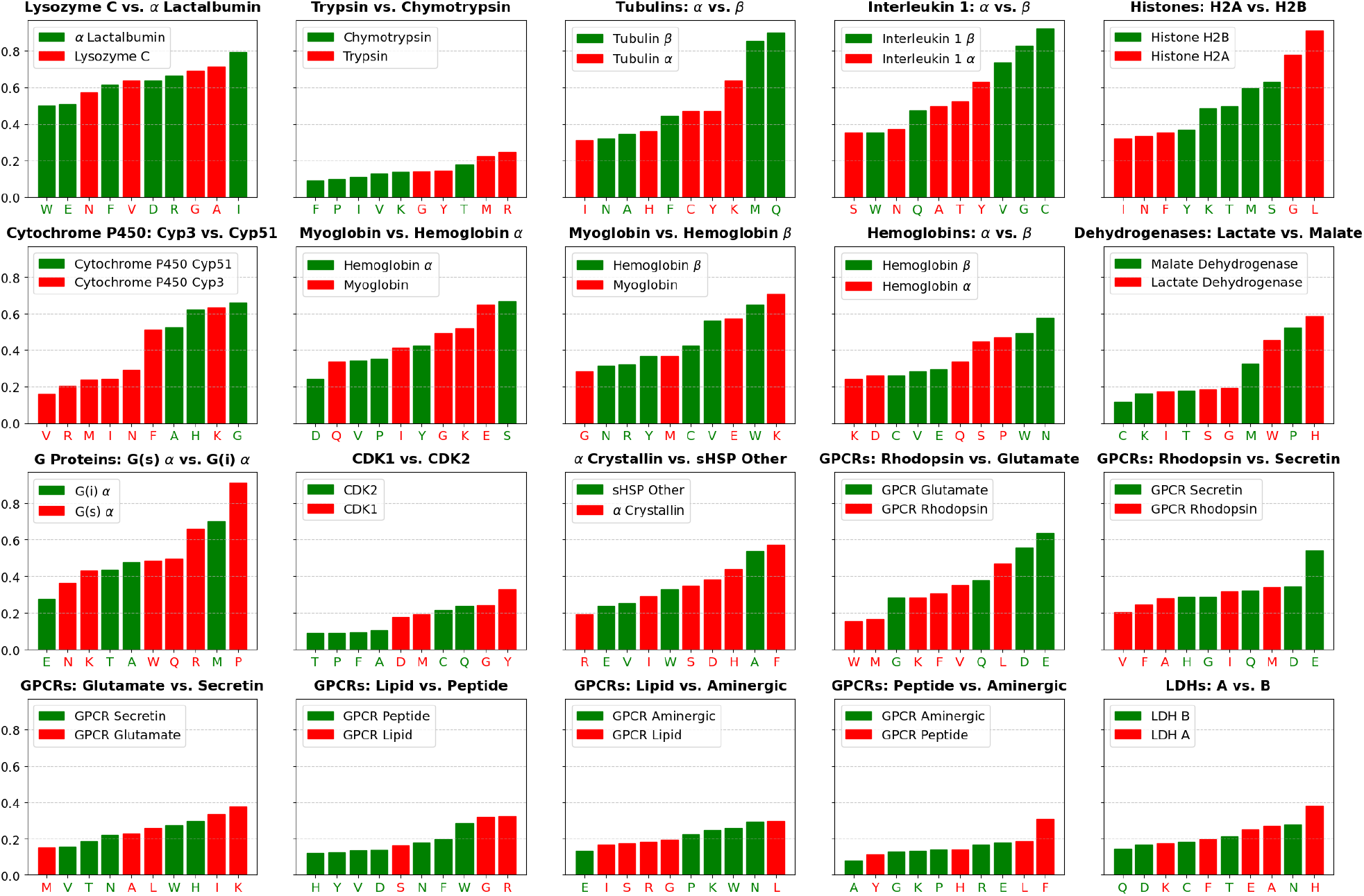
Top-10 AAC features ranked by MCI across paralog pairs. The amino acids are partitioned class-wise based on the sign (+/−) of the corresponding coefficient of the linear SVM classifier trained using the AAC features. We observe that for paralog pairs with top MCI score < 0.4, the test AM scores with full AAC features are < 0.9. A significant jump in test AM scores is observed for these pairs with 2-OAAC, as can be seen in Table 2.

### 3.2. Role of high MCI AAC features in distinguishing paralogs

We look at the top-10 AAC features (amino acids) ranked by MCI for each paralog pair, as shown in Fig. 3. Furthermore, we partition these top-10 features (amino acids) class-wise based on the sign (+/−) of the coefficients of the linear SVM classifier trained using the AAC features. For a paralog pair *P* vs *Q*, we will refer to the class-wise partition of top-10 features as *AFS*(*P*) and *AFS*(*Q*), where *AFS* means amino acid feature subset. Below, we discuss some evidence from the literature or structural analysis that highlights the role of the *AFS* amino acids in the functional difference of some of the paralog pairs in our dataset.

#### 3.2.1. Tubulins: α and β

These two paralogs have evolved to co-assemble into heterodimers, which then polymerise to form the microtubule lattice.^13^ We looked at the contact residues of a tubulin-*α* chain and a tubulin-*β* chain in the 3D structures of a microtubule lattice made of 6 tubulin-*α*/*β* heterodimers (PDB IDs: 3JAR, 5N5N). We see that the contact points of the tubulin-*α* chain in the heterodimer have more *AFS*(Tubulin-*α*) amino acids than *AFS*(Tubulin-*β*).

Similarly, *AFS*(Tubulin-*β*) amino acids are more than *AFS*(Tubulin-*α*) at the contact point of the tubulin-*β* chain in the heterodimer. Thus, the amino acids identified in *AFS* can be considered to be significant towards the quaternary structure of tubulin-*α*/*β* heterodimer.

#### 3.2.2. Histones: H2A and H2B

Histones have a heterooctameric structure comprising of two H2A/H2B dimers and one H3/H4 tetramer.^14^ We looked at the contact residues of an H2A chain and H2B chain in the heteroocatmer structure of histone (PDB IDs: 3KWQ, 1AOI). We find that the contact points of H2A chain in the heterooctamer have more *AFS*(Histone H2A) amino acids than *AFS*(Histone H2B). Thus, the amino acids identified in *AFS* can be considered to be significant towards the quaternary structure of the histone heterooctamer.

#### 3.2.3. Interleukin-1: α and Interleukin-1 β

C has the highest MCI score and is in *AFS*(Interleukin-1 *β*). Deleting C results in loss of activity in Interleukin-1 *β*.^15^ We do not find such studies for Interleukin-1 *α*.

#### 3.2.4. Cytochrome P450: CYP3 and CYP51

H and G with the highest MCI score in *AFS*pCYP51q have been reported^16−19^ to be important in the enzymatic activity of CYP51. Mutation of these amino acids at specific positions has been shown to result in a decrease in the activity of the enzyme.^17,18^ Similarly, F with the second-highest MCI in *AFS*pCYP3q is also known to be important in the enzymatic activity of CYP3.^20−22^ A cluster of F residues in CYP3 is known to form a substrate-binding pocket with an active site.^22^

#### 3.2.4. Globins

Myoglobin is a monomer, while *α* and *β* chains together constitute hemoglobin, a tetramer of composition *α*_2_*β*_2_.^23^ We superimposed the 3D structures of myoglobin, hemoglobin-*α* and hemoglobin-*β* (PDB IDs: 3RGK, 1HHO) and mapped the *α, β* contact residues (based on Ref. 24) of hemoglobin tetramer to that of myoglobin. We want to check if the respective *AFS* amino acids have a role in the difference in the oligomerisation of myoglobin and hemoglobin.

K and E are the top-ranked amino acids in *AFS*(Myoglobin) of both ‘Myoglobin vs Hemoglobin-*α*’ and ‘Myoglobin vs Hemoglobin-*β*’. We find that the amino acids K, E are less in number at the contact residues of hemoglobin tetramer and more in number at the corresponding locations in myoglobin, which is a monomer (see Fig. 4).

**Fig. 4.**
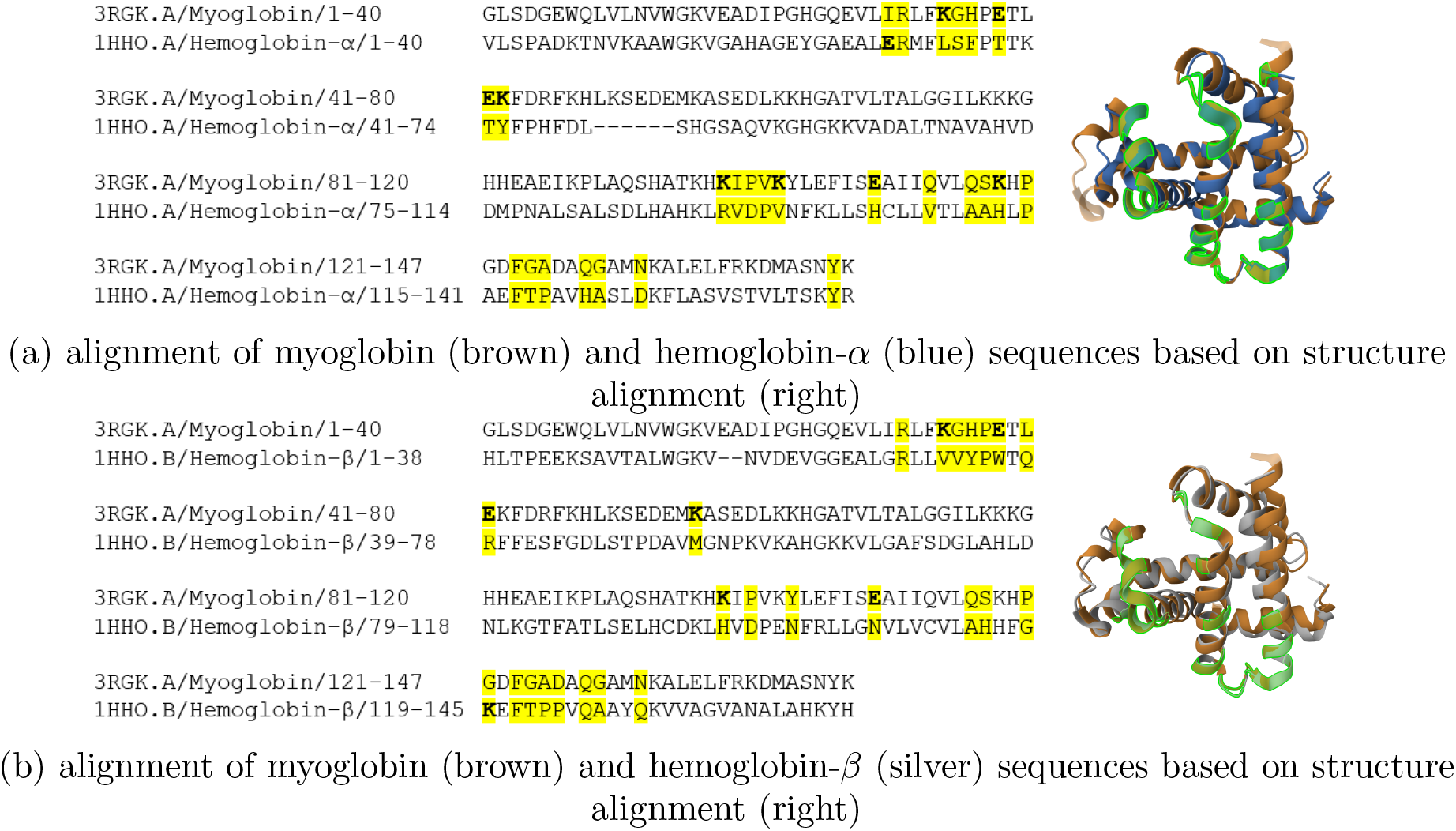
Structural alignment of myoglobin with hemoglobin-*α* (a) and hemoglobin-*β* (b). Hemoglobin tetramer contact points (Ref. 24) are highlighted in yellow in the sequences and green in the 3D structures (right). The corresponding aligned positions in the myoglobin sequence are similarly highlighted. Notably, amino acids K and E emerged as the top distinguishing features for myoglobin in both independent comparisons (myoglobin vs hemoglobin-*α* and myoglobin vs hemoglobin-*β*). As seen here, these two amino acids appear significantly more frequently at these loci in monomeric myoglobin than in the tetrameric hemoglobin contacts. Structure alignments were generated via the RCSB PDB tool: https://www.rcsb.org/alignment (jFATCAT-rigid algorithm, default parameters).

##### Hemoglobins

*α and β*. We looked at the contact residues of an *α* and *β* chain in the tertrameric hemoglobin structure (PDB IDs: 3KWQ, 1A3N). We find that the *AFS*(Hemoglobin-*α*) amino acids are more in the *α* chain contact points than the *β* chain contact points. Similarly, the other way holds for *AFS*(Hemoglobin-*β*).

## 4. Discussion and Conclusions

### AAC features have good predictive performance in distinguishing paralogs

Despite the lack of sequence order information, AAC proves surprisingly powerful at distinguishing paralog pairs. For many of the pairs in our dataset (13 of 20), AAC alone yielded test AM scores exceeding 0.9. A recent review study, Ref. 25, highlights that the amino acid composition of a protein can sometimes be the key driver of its activity. The ability to separate paralog sequences linearly solely based on AAC strongly backs this concept, indicating composition-driven signatures associated with their specialised biological roles.

### MCI scores of AAC features can point to amino acids that have a potential role in the functional differences of parlogs

Using the MCI framework on our paralog pair datasets, we illustrate a robust, theoretically grounded method for identifying specific amino acids that potentially drive the paralog divergence. Crucially, using only the top-*k* amino acids ranked by MCI retained nearly all of the predictive power and, in some cases, acted as a regulariser to improve generalisation. Furthermore, our structural mapping analysis (e.g., Tubulins, Histones, Globins) provided biological validation: the amino acids flagged by high MCI scores frequently map to critical functional loci, such as protein-protein interaction interfaces responsible for specific oligomerisation states (e.g., the monomeric state of myoglobin vs. the tetrameric state of hemoglobin). This illustrates that MCI can be a viable tool for generating hypotheses about protein functions.

### The minimal sequence order information of 2-OAAC features enables very high predictive performance in distinguishing paralog pairs

While AAC is highly discriminatory for most pairs, it struggles with certain challenging paralogs (e.g., LDH-A vs. LDH-B or Trypsin vs. Chymotrypsin), where the functional differences are nuanced. In these cases, we see that introducing minimal sequence order information via 2-OAAC significantly boosts predictive performance. A second-order feature space, which tracks pairs of residues rather than singletons, represents the absolute minimum amount of sequence order information one can incorporate beyond basic composition. 2-OAAC is one such minimal representation. 2-OAAC captures the relative occurrences of all ordered amino acid pairs across the entire protein length, regardless of their separation distance. This allows it to encode long-range ordered compositional asymmetries along the sequence that composition alone cannot fully capture. The near-perfect linear separation achieved by 2-OAAC across almost all paralog pairs underscores the importance of capturing the ordered compositional asymmetry in the sequence for distinguishing highly similar paralog pairs.

Furthermore, we also evaluated third-order sequence context using 3-OAAC features (see results in Table S1 of Supplementary File 1). While it resulted in marginal performance improvements for most families, the test AM scores dropped significantly for challenging pairs like LDH-A vs LDH-B due to overfitting on exponentially high-dimensional feature space.

### The challenge of low sample-size and label noise

To ensure we had sufficient samples for training and testing, our dataset inclusion criteria allowed for sequences with lower-quality, computationally-derived annotations (e.g., Unreviewed/TrEMBL entries). As a result, the datasets inevitably contain electronically annotated sequences, introducing potential label noise and even out-of-distribution samples. This is particularly relevant for pairs such as Trypsin/Chymotrypsin and LDH-A/LDH-B, where we had to rely heavily on such annotations to bolster sample sizes (as reflected in the broad database queries detailed in Supplementary File 1). Consequently, the relatively lower classification test scores observed for these specific families (especially LDH-A/LDH-B) could partially stem from this label noise and low sample size rather than an inherent lack of biological separability. Estimating the exact extent of this noise computationally is challenging and would ultimately require wet-lab experimentation to correct.

### Future directions

Our results open several new directions for study. In this study, to manage the high dimensionality of 2-OAAC features, we devised an amino acid-wise MCI (aa-wise MCI). A direct extension of this work would be to conduct a deeper biological and structural analysis of the top-ranked amino acids by this aa-wise MCI for 2-OAAC. Furthermore, one could investigate the exact arg max of the aa-wise MCI to pinpoint the specific ordered amino acid pairs that carry high marginal contributions. Biologically, these highly discriminative amino acids and sequence-order pairs provide prime candidates for targeted mutagenesis experiments, offering an accelerated computational method to guide the experimental validation of paralog-specific functions. Another promising direction for future work is label noise robust binary classification methods, to address the inherent label noise in biological datasets.

## References

1. The UniProt Consortium, UniProt: the universal protein knowledgebase in 2021, Nucleic Acids Research 49, D480 (11 2020).

2. Y. Yang, T. Xu, G. Conant, H. Kishino, J. L. Thorne and X. Ji, Interlocus gene conversion, natural selection, and paralog homogenization, Molecular Biology and Evolution 40, p. msad198 (09 2023).

3. B. Rost, Twilight zone of protein sequence alignments, Protein Engineering 12, 85 (02 1999).

4. N. Kresge, R. D. Simoni and R. L. Hill, The development of site-directed mutagenesis by michael smith, Journal of Biological Chemistry 281, e31 (2006).

5. A. Catav, B. Fu, Y. Zoabi, A. L. W. Meilik, N. Shomron, J. Ernst, S. Sankararaman and R. Gilad-Bachrach, Marginal contribution feature importance - an axiomatic approach for explaining data, in Proceedings of the 38th International Conference on Machine Learning, eds. M. Meila and T. Zhang, Proceedings of Machine Learning Research, Vol. 139 (PMLR, 18–24 Jul 2021).

6. M. O. Dayhoff, R. M. Schwartz and B. C. Orcutt, A model of evolutionary change in proteins, Atlas of protein sequence and structure 5, 89 (1972).

7. J. Shen, J. Zhang, X. Luo, W. Zhu, K. Yu, K. Chen, Y. Li and H. Jiang, Predicting protein–protein interactions based only on sequences information, Proceedings of the National Academy of Sciences 104, 4337 (2007).

8. W. R. Taylor, The classification of amino acid conservation, Journal of Theoretical Biology 119, 205 (1986).

9. A. K. Menon, H. Narasimhan, S. Agarwal and S. Chawla, On the statistical consistency of algorithms for binary classification under class imbalance, in Proceedings of the 30th ICML - Volume 28, (JMLR.org, 2013).

10. K. H. Brodersen, C. S. Ong, K. E. Stephan and J. M. Buhmann, The Balanced Accuracy and Its Posterior Distribution, in Pattern Recognition, International Conference on, (IEEE Computer Society, Los Alamitos, CA, USA, August 2010).

11. S. Tripathi, N. Hemachandra and P. Trivedi, Interpretable feature subset selection: A Shapley value based approach, in 2020 IEEE BigData, (IEEE Xplore, 2020).

12. J. Castro, D. Gómez and J. Tejada, Polynomial calculation of the shapley value based on sampling, Computers & Operations Research 36, 1726 (2009), Selected papers presented at the Tenth International Symposium on Locational Decisions (ISOLDE X).

13. T. Mühlethaler, D. Gioia, A. E. Prota, M. E. Sharpe, A. Cavalli and M. O. Steinmetz, Comprehensive analysis of binding sites in tubulin, Angewandte Chemie International Edition 60, 13331 (2021).

14. S. Dutta, I. V. Akey, C. Dingwall, K. L. Hartman, T. Laue, R. T. Nolte, J. F. Head and C. W. Akey, The crystal structure of nucleoplasmin-core: Implications for histone binding and nucleosome assembly, Molecular Cell 8, 841 (2001).

15. B. Veerapandian, G. L. Gilliland, R. Raag, A. L. Svensson, Y. Masui, Y. Hirai and T. L. Poulos, Functional implications of interleukin-1β based on the three-dimensional structure, Proteins: Structure, Function, and Bioinformatics 12, 10 (1992).

16. Y. Nitahara, K. Kishimoto, Y. Yabusaki, O. Gotoh, Y. Yoshida, T. Horiuchi and Y. Aoyama, The amino acid residues affecting the activity and azole susceptibility of rat cyp51 (sterol 14-demethylase p450), The Journal of Biochemistry 129, 761 (2001).

17. G. I. Lepesheva and M. R. Waterman, Cyp51-the omnipotent p450, Molecular and Cellular Endocrinology 215, 165 (2004), Proceedings of the Serono Foundation for the Advancement of Medical Science Workshop on Molecular Steroidogenesis.

18. G. I. Lepesheva and M. R. Waterman, Sterol 14α-demethylase cytochrome p450 (cyp51), a p450 in all biological kingdoms, Biochimica et Biophysica Acta (BBA) - General Subjects 1770, 467 (2007), P450.

19. N. Strushkevich, S. A. Usanov and H.-W. Park, Structural basis of human cyp51 inhibition by antifungal azoles, Journal of Molecular Biology 397, 1067 (2010).

20. H. Qiu, S. Taudien, H. Herlyn, J. Schmitz, Y. Zhou, G. Chen, R. Roberto, M. Rocchi, M. Platzer and L. Wojnowski, Cyp3 phylogenomics: evidence for positive selection of cyp3a4 and cyp3a7, Pharmacogenetics and Genomics 18, p. 53—66 (January 2008).

21. I. G. Denisov, Y. V. Grinkova, P. Nandigrami, M. Shekhar, E. Tajkhorshid and S. G. Sligar, Allosteric interactions in human cytochrome p450 cyp3a4: The role of phenylalanine 213, Biochemistry 58, 1411 (2019), PMID: 30785734.

22. Y. Zhang, Z. Wang, Y. Wang, W. Jin, Z. Zhang, L. Jin, J. Qian and L. Zheng, Cyp3a4 and cyp3a5: the crucial roles in clinical drug metabolism and the significant implications of genetic polymorphisms, PeerJ 12, p. e18636 (2024).

23. K. Dill, R. Jernigan and I. Bahar, Protein Actions: Principles and Modeling (CRC Press, 2017).

24. M. Shionyu, K. Takahashi and M. Gō, Variable subunit contact and cooperativity of hemoglobins, J. Mol. Evol. 53, 416 (October 2001).

25. S. M. Cascarina and E. D. Ross, Protein activities driven by amino acid composition, Journal of Biological Chemistry 301, p. 110640 (2025).

