## Supplementary File 1 for "Alignment-free sequence composition based features can distinguish paralogous proteins"

**Supplementary file 1 for the paper titled: *Alignment-free sequence composition based features can distinguish paralogous proteins***

### S1. Dataset details

#### S1.1. *Data filtering and preprocessing*

To ensure high-quality and consistent datasets for classification, the following automated filtering steps were applied to the raw sequences downloaded from the UniProt database for each protein family:

(1) **Baseline length calculation:** We first extracted the manually reviewed (SwissProt) sequences and calculated the mean ($\mu$) and standard deviation ($\sigma$) of their sequence lengths.
(2) **Length-based outlier removal:** We established acceptable sequence length bounds using the 3-sigma rule ($\mu \pm 3\sigma$). After combining the SwissProt and the unreviewed (TrEMBL) sequences, any sequence falling outside these bounds was discarded. This effectively removes extremely short fragments and abnormally long fusion proteins.
(3) **LCSS-based redundancy filtering:** To reduce bias and ensure a challenging dataset, we performed a sequence redundancy check using the longest common subsequence score (LCSS). We define the LCSS between two sequences as the length of their longest common subsequence divided by the length of the longer of the two sequences. We computed pairwise LCSS for all sequences in a family and constructed a graph connecting sequences with an LCSS $\geqslant 0.95$. We then extracted connected components and retained only a single representative sequence per component, ensuring no two sequences in the final dataset share $\geqslant 95\%$ subsequence similarity.

Following these filtering steps, the resulting quality-controlled sequences for each family were randomly split into 80% training and 20% test sets.

#### S1.2. *UniProt search queries*

The following specific query strings were used via the UniProt REST API to fetch the sequences for each paralog family. All queries explicitly excluded fragments (`fragment:false`) and sequences with low existence/annotation scores (`existence:4, existence:5`).

- **Lysozyme-like:**
  - *Lysozyme C*: `(protein_name:"lysozyme C") AND (ec:3.2.1.17)`
  - *α-Lactalbumin*: `((protein_name:"alpha lactalbumin") OR (protein_name:"Lactose synthase B protein"))`
- **Trypsin-like protease:**
  - *Trypsin*: `(family:"peptidase S1 family") AND (protein_name:trypsin) AND (ec:3.4.21.4)`
  - *Chymotrypsin*: `(family:"peptidase S1 family") AND (protein_name:chymotrypsin) AND (ec:3.4.21.1)`

- **Tubulin:**
  - *Tubulin $\alpha$*: (protein_name:"tubulin alpha") AND (family:"tubulin family")
  - *Tubulin $\beta$*: (protein_name:"tubulin beta") AND (family:"tubulin family")
- **Interleukin-1 (IL-1):**
  - *IL-1 $\alpha$*: (gene:IL1A) AND (family:il-1)
  - *IL-1 $\beta$*: (gene:IL1B) AND (family:il-1)
- **Histone:**
  - *H2A*: (protein_name:"histone h2a") AND (family:histone)
  - *H2B*: (protein_name:"histone h2b") AND (family:histone)
- **Cytochrome P450:**
  - *Cyp3*: (family:"Cytochrome P450") AND ((gene:cyp3) OR (gene:cyp3A*))
  - *Cyp51*: (family:"Cytochrome P450") AND ((gene:cyp51) OR (gene:cyp51A*) OR (gene:cyp51B*) OR (gene:cyp51C*))
- **Globins:**
  - *Myoglobin*: (protein_name:"myoglobin") AND (xref:interpro-IPR002335)
  - *Hemoglobin-$\alpha$*: (protein_name:"hemoglobin alpha") AND (xref:interpro-IPR002338)
  - *Hemoglobin-$\beta$*: (protein_name:"hemoglobin beta") AND (xref:interpro-IPR002337)
- **G-Protein Coupled Receptors (GPCRs):**
  - *Rhodopsin-like*: (family:"g-protein coupled receptor 1 family")
  - *Lipid*: (family:"g-protein coupled receptor 1 family") AND ((protein_name:cannabinoid) OR ... OR (protein_name:oxylipin))
  - *Peptide*: (family:"g-protein coupled receptor 1 family") AND ((protein_name:angiotensin) OR ... OR (protein_name:cholecystokinin))
  - *Aminergic*: (family:"g-protein coupled receptor 1 family") AND ((protein_name:dopamine) OR ... OR (protein_name:"trace amine"))
  - *Glutamate-like*: (family:"g-protein coupled receptor 3 family")
  - *Secretin-like*: (family:"g-protein coupled receptor 2 family")
- **Dehydrogenase:**
  - *Lactate Dehydrogenase (LDH)*: (protein_name:"lactate dehydrogenase") AND (family:"ldh mdh superfamily")
  - *Malate Dehydrogenase (MDH)*: (protein_name:"malate dehydrogenase") AND (family:"ldh mdh superfamily")
- **LDH Subtypes:**
  - *LDH-A*: (gene:ldha) AND (family:"ldh mdh superfamily") NOT (gene:ldhb)
  - *LDH-B*: (gene:ldhb) AND (family:"ldh mdh superfamily") NOT (gene:ldha)
- **G(alpha) Subunits:**

- *G(s) α*: `(protein_name:"G(s) subunit alpha") AND (family:"g-alpha family")`
- *G(i) α*: `(protein_name:"G(i) subunit alpha") AND (family:"g-alpha family")`

- **Cyclin-dependent Kinases (CDK):**
  - *CDK1*: `(protein_name:"cyclin-dependent kinase 1") AND (family:"protein kinase superfamily")`
  - *CDK2*: `(protein_name:"cyclin-dependent kinase 2") AND (family:"protein kinase superfamily")`
- **Small Heat Shock Proteins (sHSP):**
  - *α-Crystallin*: `(protein_name:"alpha-crystallin") AND (family:"small heat shock protein (hsp20) family")`
  - *Other sHSP*: `(family:"small heat shock protein (hsp20) family") NOT (protein_name:"alpha-crystallin")`

### S2. Supplementary Results

Table S1. Classification performance (AM scores) across paralog pairs using 3-OAAC features.

| Paralog Pair | Train AM | Val AM | Test AM |
|---|---|---|---|
| Lysozyme C vs $\alpha$-Lactalbumin | 0.99 | 0.98 | 1.00 |
| Trypsin vs Chymotrypsin | 0.94 | 0.89 | 0.90 |
| Tubulins: $\alpha$ vs $\beta$ | 0.98 | 0.98 | 1.00 |
| Interleukin-1: $\alpha$ vs $\beta$ | 1.00 | 1.00 | 1.00 |
| Histones: H2A vs H2B | 1.00 | 1.00 | 0.98 |
| Cytochrome P450: Cyp3 vs Cyp51 | 0.98 | 0.98 | 0.98 |
| Myoglobin vs Hemoglobin $\alpha$ | 1.00 | 1.00 | 0.99 |
| Myoglobin vs Hemoglobin $\beta$ | 1.00 | 1.00 | 1.00 |
| Hemoglobins: $\alpha$ vs $\beta$ | 1.00 | 0.99 | 1.00 |
| Dehydrogenases: Lactate vs Malate | 0.99 | 0.98 | 0.99 |
| LDHs: A vs B | 0.78 | 0.69 | 0.57 |
| G Proteins: G(s) $\alpha$ vs G(i) $\alpha$ | 1.00 | 1.00 | 1.00 |
| CDK1 vs CDK2 | 0.93 | 0.92 | 0.94 |
| $\alpha$-Crystallin vs sHSP Other | 0.97 | 0.97 | 0.97 |
| GPCRs: Rhodopsin vs Glutamate | 1.00 | 0.99 | 0.98 |
| GPCRs: Rhodopsin vs Secretin | 0.99 | 0.99 | 0.99 |
| GPCRs: Glutamate vs Secretin | 0.98 | 0.97 | 0.95 |
| GPCRs: Lipid vs Peptide | 0.96 | 0.94 | 0.94 |
| GPCRs: Lipid vs Aminergic | 1.00 | 0.99 | 1.00 |
| GPCRs: Peptide vs Aminergic | 0.99 | 0.98 | 0.98 |
